# Characterization of Genome-wide Phylogenetic Conflict Uncovers Evolutionary Modes of Carnivorous Fungi

**DOI:** 10.1101/2024.03.21.586083

**Authors:** Weiwei Zhang, Yani Fan, Wei Deng, Yue Chen, Shunxian Wang, Seogchan Kang, Jacob Lucas Steenwyk, Meichun Xiang, Xingzhong Liu

**Affiliations:** State Key Laboratory of Medicinal Chemical Biology, Key Laboratory of Molecular Microbiology and Technology, Department of Microbiology, College of Life Science, Nankai University, Tianjin 300071, China; State Key Laboratory of Mycology, Institute of Microbiology, Chinese Academy of Sciences, Beijing 100101, China; University of Chinese Academy of Sciences, Beijing 100049, China; Department of Plant Pathology & Environmental Microbiology, The Pennsylvania State University, PA 16802, USA; Howards Hughes Medical Institute and Department of Molecular and Cell Biology, University of California, Berkeley, CA 94720, USA

## Abstract

Mass extinction has often paved the way for rapid evolutionary radiation, resulting in the emergence of diverse taxa within specific lineages. While the emergence and diversification of carnivorous nematode-trapping fungi (NTF) in Ascomycota has been linked to the Permian-Triassic (PT) extinction, the processes underlying NTF radiation remain unclear. Here, we conducted phylogenomic analyses using 23 genomes spanning three NTF lineages, each employing distinct nematode traps — mechanical traps (*Drechslerella* spp.), three-dimensional (3-D) adhesive traps (*Arthrobotrys* spp.), and two-dimensional (2-D) adhesive traps (*Dactylellina* spp.), and one non-NTF species as the outgroup. This analysis revealed how diverse mechanisms contributed to the tempo of NTF evolution and rapid radiation. The genome-scale species tree of NTFs suggested that *Drechslerella* emerged earlier than *Arthrobotrys* and *Dactylellina*. Extensive genome-wide phylogenetic discordance was observed, mainly due to incomplete lineage sorting (ILS) between lineages (∼81.3%). Modes of non-vertical evolution (i.e., introgression and horizontal gene transfer) also contributed to phylogenetic discordance. The ILS genes that are associated with hyphal growth and trap morphogenesis (e.g., those associated with the cell membrane system and cellular polarity division) exhibited signs of positive selection.

## Introduction

Mass extinctions result in vacated ecological niches that can be occupied by novel species and drive subsequent radiation events (Sepkoski 1998; Jablonski 2001). Mass extinction and concomitant radiations have been documented in multiple lineages, including angiosperms (Silvestro et al. 2015), planktic foraminifera (Lowery and Fraass 2019), snakes (Grundler and Rabosky 2021), modern birds (Jarvis et al. 2014), and mushrooms (Varga et al., 2019). Comparative genomics has facilitated systematic identification of candidate genetic changes underlying speciation and adaptive radiation (Marques et al. 2019).

Carnivorous nematode-trapping fungi (NTF) emerged after the Permian-Triassic (PT) extinction and radiated into multiple lineages that form distinct trapping devices to capture free-living nematodes (Yang et al. 2012). The emergence of NTF from saprophytic fungal species is thought to be driven by nematode proliferation that occurred in the wake of the PT extinction, an event that resulted in a carbon-rich and nitrogen-poor environment (Barron, 1977; Gray, 1983; Liu et al. 2014; Fan et al. 2024). The ability to capture and consume nematodes allows NTF to obtain extra nitrogen, likely conferring a competitive advantage over saprophytic fungi (Yang et al. 2012) and driving diversification. NTF have radiated into three clades, each with unique genus designations and trapping systems—*Arthrobotrys* spp. employ three-dimensional (3D) adhesive traps (networks); *Dactylellina* spp. utilize two-dimensional (2D) adhesive traps (knob, column, non-constricting ring); and *Drechslerella* spp. form mechanical traps (constricting ring) (Jiang et al., 2017).

Phylogenomics has greatly advanced our understanding of the Tree of Life, mechanisms of gene and genome evolution, and relationship between genomic and phenotypic divergence during speciation (Jin et al. 2021; Steenwyk et al. 2023). Phylogenomics has also revealed how individual genes have undergone evolutionary histories that are distinct from the phylogenetic history of species carrying these genes (Steenwyk et al. 2019; Salichos and Rokas 2013). Theoretical and empirical studies have shown that this discordance or incongruence can be caused by analytical factors, such as errors in taxon sampling and gene tree estimation, and biological mechanisms, such as incomplete lineage sorting (ILS), horizontal gene transfer (HGT), and introgressive hybridization (Steenwyk et al. 2020a; Lopes et al. 2021; Shen et al. 2021; Feng et al. 2022; Steenwyk et al. 2023). Other factors also influenced species diversification during radiation. For example, adaptive evolution punctuated by positive selection occurs more frequently in radiating lineages than in slowly diversifying ones (Nevado et al. 2019). While ILS, HGT, introgression, and positive selection have been documented in several eukaryotic lineages, such as cichlids (Brawand et al. 2014), wild tomatoes (Pease et al. 2016), honeybees (Fouks et al. 2021), big cats (Figueiró et al. 2017), and *Populus* species (Wang et al. 2020), their impact on fungal radiation events remains poorly understood.

Here, we characterized the genome-wide patterns and drivers of phylogenetic discordance in the three NTF lineages. Patterns of genome-wide phylogenetic discordance showed that ILS between lineages caused most of the observed discordances. In contrast, introgression and HGT contributed less to the incongruence between the species tree and gene trees. Positive selection of ILS genes associated with growth and trap morphogenesis were also observed. Similar to previous studies of other lineages, our phylogenomic analyses revealed how diverse evolutionary mechanisms contributed to the tempo of NTF evolution and rapid radiation.

## Results

### Extensive phylogenomic discordance among NTF

To investigate the evolutionary history of NTF in Ascomycota, we analyzed 23 NTF genomes (Supplementary Table S1). Our taxon sampling covered three major lineages that underwent radiation and evolved distinct mechanisms of nematode trapping, including 3-D adhesive networks (*Arthrobotrys* spp.), 2-D adhesive traps (*Dactylellina* spp.), and mechanical traps (*Drechslerella* spp.), and *Dactylella cylindrospora*, a non-NTF closely related to NTF as the outgroup.

Single-copy orthologous genes (2,944 in total; Supplementary Table S2) present in all species were combined to construct maximum likelihood species tree using two alignment and trimming strategies (Clustal-Omega + ClipKIT and MAFFT + Gblocks) (Castresana 2000; Katoh and Standley 2013; Sievers and Higgins 2018; Steenwyk et al. 2020b). The species tree topologies were consistent under both strategies (Figure 1, Figure S1), suggesting our analyses are robust to some analytical sources of error associated with alignment and trimming strategy (Steenwyk et al. 2023). The genome-scale phylogeny was consistent with our previously published multiple-gene phylogeny (Yang et al. 2007) and strongly supported the placement of *Arthrobotrys* and *Dactylellina* as sister genera (Figure 1). The species tree supported two notable evolutionary events: the divergence of those forming mechanical traps (*Drechslerella*) and the lineage that produces adhesive traps and the subsequent divergence of 2-D (*Dactylellina*) and 3-D (*Arthrobotrys*) adhesive traps.

**Figure 1.**
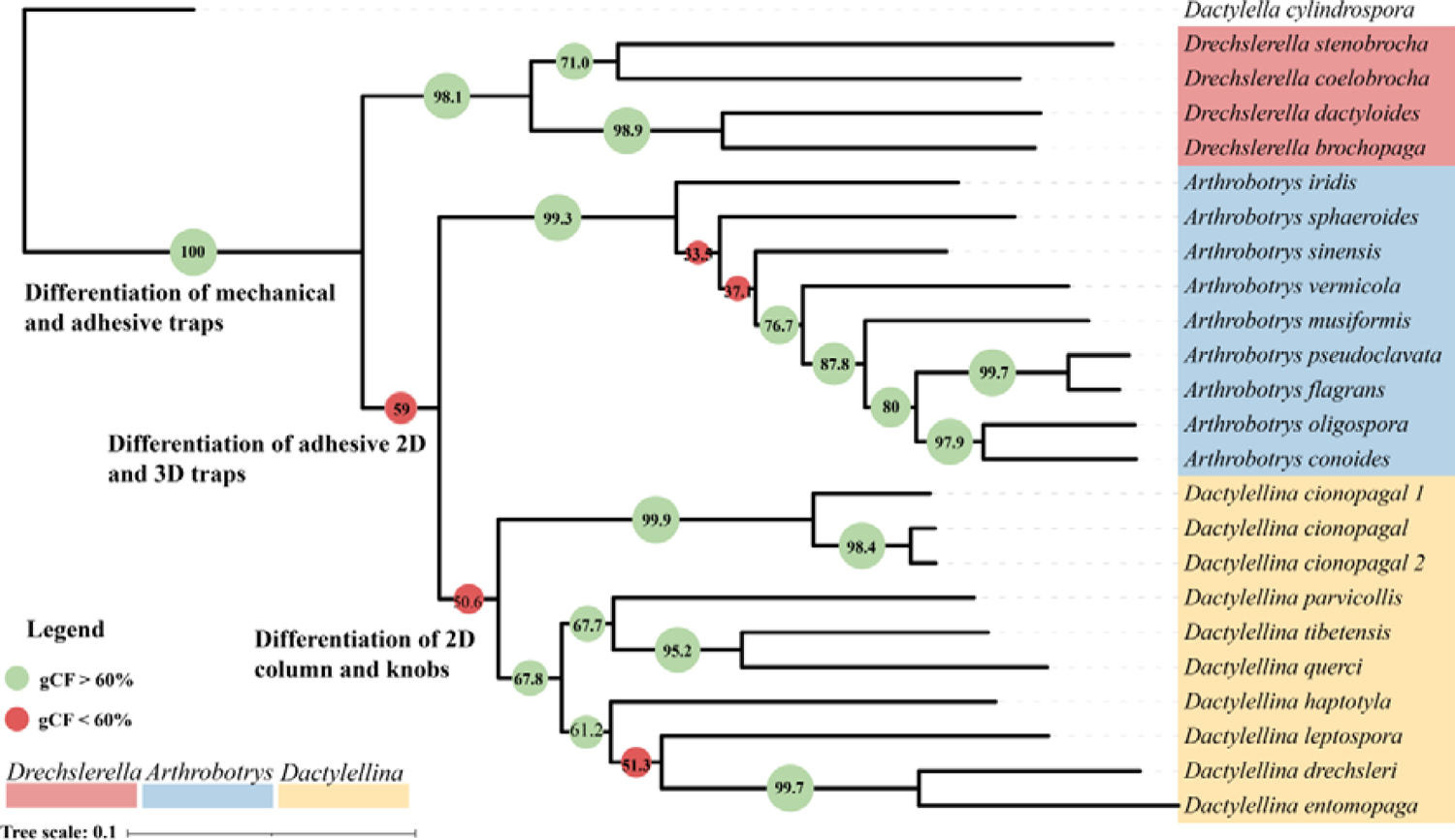
Phylogeny of nematode-trapping fungi. Their phylogenic relationships were determined using concatenated nucleotide sequences of the single-copy orthologous genes present in all species. *Dactylella cylindrospora*, a non-NTF species, was used as the outgroup. Bootstrap values were 100% on each node. Gene-concordance factors (gCF) values were calculated by IQ-TREE and annotated on each node, with green indicating nodes greater than 60% and red indicating nodes less than 60%.

Maximum likelihood trees of single-copy orthologous genes were also constructed using Clustal-Omega + ClipKIT and MAFFT + Gblocks approaches. The resulting trees were largely consistent between each method, suggesting analytical errors associated with software choice are minimal (Steenwyk et al. 2023). Nonetheless, discordance between the single-gene trees and the species tree was abundant (Figure 2, Supplementary Table S2). Densitree plots depicted numerous topological conflicts among the gene trees (Figure 2a), and MDS analysis based on Robinson-Foulds (RF) distances revealed differences between the gene trees and the species tree (Figure 2b). Concordance analyses based on IQ-TREE showed that there was a high rate of conflict between gene trees and the species tree at the divergence points between mechanical traps (*Drechslerella*) and adhesive traps, as well as between 2-D (*Dactylellina*) and 3-D (*Arthrobotrys*) adhesive traps (gene-concordance factors (gCF) < 60%, Figure 1). Furthermore, there were two and one nodes with high conflict (gCF < 60%, Figure 1) within *Arthrobotrys* and *Dactylellina*, respectively.

**Figure 2.**
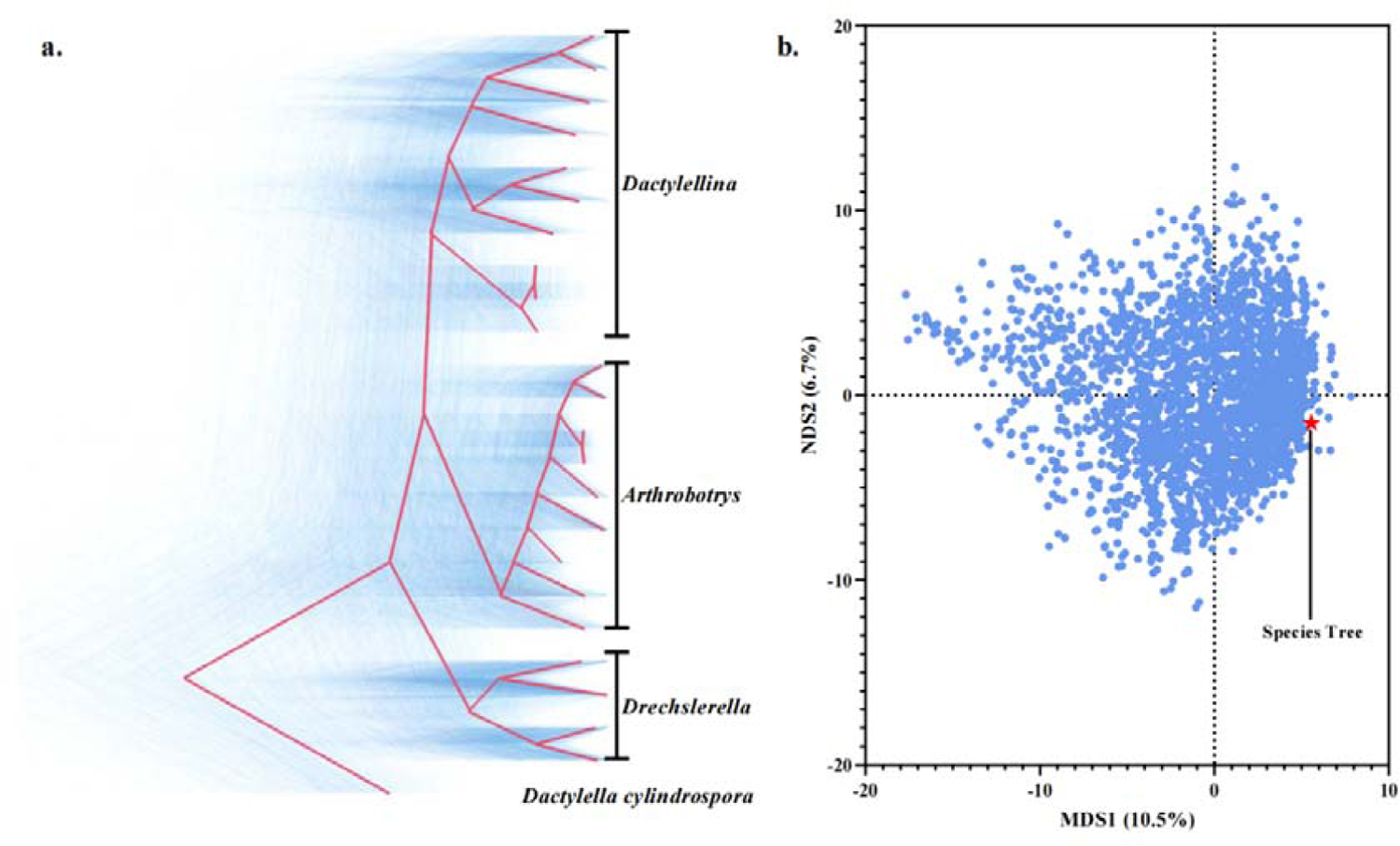
Extensive conflict between the gene trees and the species tree. a. Densitree plot. Blue represents the gene trees, and red represents the consensus tree inferred by the Densitree software, which is consistent with the topology of the species tree. b. A plot resulting from multi-dimensional scaling (MDS) analysis illustrates the topological differences between the gene trees (denoted by blue dots) and the species tree (denoted by the red pentagram).

### ILS is largely responsible for phylogenetic discordance

To further rule out analytical sources of error, we identified single gene trees that were consistent between the two alignment and trimming strategies — Clustal-Omega + ClipKIT and MAFFT+Gblocks (Castresana 2000; Katoh and Standley 2013; Sievers and Higgins 2018; Steenwyk et al. 2020b). Among the 2,944 single-copy orthologous genes, 64 orthologous genes yielded inconsistent gene trees with the two strategies (see Supplementary Table S2); inconsistent genes, which are likely subject to analytical errors, were removed from subsequent analyses.

Among the remaining 2,880 gene trees, 496 exhibited average bootstrap support below 80% (Supplementary Table S1), suggesting that errors in phylogenetic inference may have affected these trees. Among the remaining 2,392 trees with high bootstrap support, 978 trees (40.9%; a group designated as Tree1) supported the species tree, whereas 1,414 trees (59.1%) were inconsistent with the species tree.

The Multispecies Coalescent (MSC) model was employed to investigate whether the observed topologies of gene trees across sets of four lineages could be attributed to ILS. By employing hypothesis testing on 1,414 gene trees, we assessed the concordance between observed gene tree distributions and those predicted under the MSC model. The results showed that at the 0.0001 significance level, 81.3% of the four-lineage scenarios supported the hypothesis that ILS shaped the topology. Whereas 18.7% of the scenarios rejected the hypothesis (Figure 3a), suggesting other evolutionary modes, like introgression and HGT, may influence the history of these loci.

**Figure 3.**
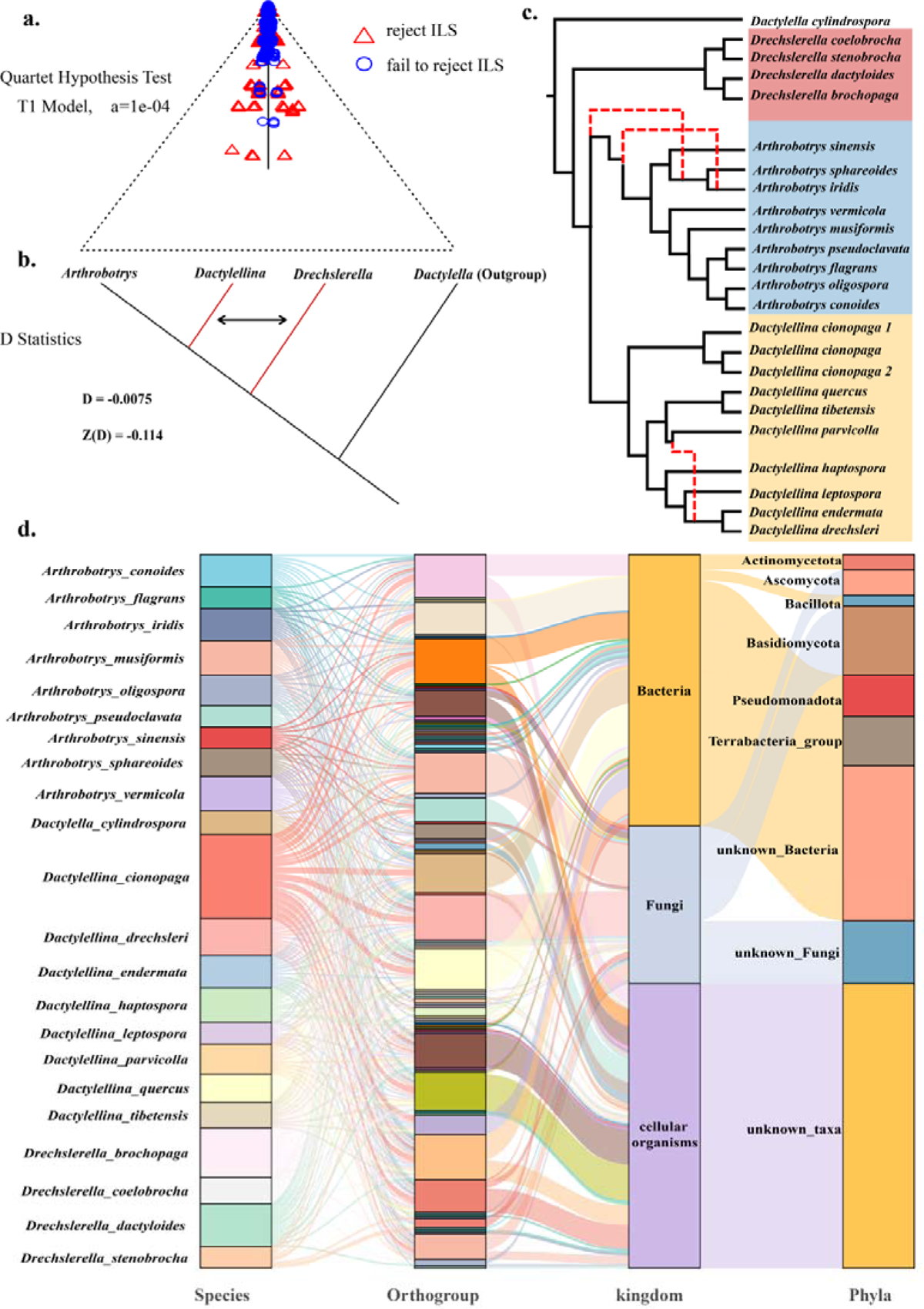
Origins of conflict between the gene trees and the species tree. a. ILS analysis based on the Multispecies coalescent (MSC) analysis. Blue circles represent four-taxa scenarios in which the topology can be explained solely by the ILS. Red triangles represent scenarios in which this hypothesis is rejected, indicating that the topology is explained by other factors. The closer the blue circles to the center of the triangle, the stronger the influence of ILS. b. Schematic representation of D-statistic results. c. Reticulate phylogenetic tree inferred by Phylonet, with red indicating gene introgression sites. When the number of hybridization events was set to 3, the tree inferred by PhyloNet matched with the species tree, and the fit was optimal. d. Sankey diagram depicting the suspected HGT events among NTF and the predicted sources of the genes.

Examination of genome-wide D-statistics analysis (also known as the ABBA-BABA test; Hibbins and Hahn, 2022; Bjornson et al. 2023), which test for introgression, revealed insignificant amounts of introgression among the three NTF lineages (Figure 3b; D = −0.0075, Z = −0.114). However, phylogenetic network analysis revealed two gene introgression events in *Arthrobotrys* lineage and one in *Dactylellina* lineage (Figure 3c); notably, these nodes high degrees of conflict among gene trees and the species tree in Figure 1.

Among the 1,414 genes displaying topological structures that conflict with the species tree, 36 appeared to have been acquired via HGT. These HGT genes predominantly originated from bacteria, with Pseudomonadota being the main donor phylum. Additionally, some HGT events from fungi, particularly from the sister phylum Basidiomycota, were also observed.

The remaining 1,378 trees were categorized into three groups (Figure 4): 7.0% (97) placed *Arthrobotrys* and *Drechslerella* as sister groups (Tree2), 18.0% (245) clustered *Drechslerella* and *Dactylellina* (Tree3), and 75.0% (1,036) did not align with their corresponding generic clades (designated as Unclassified).

**Figure 4.**
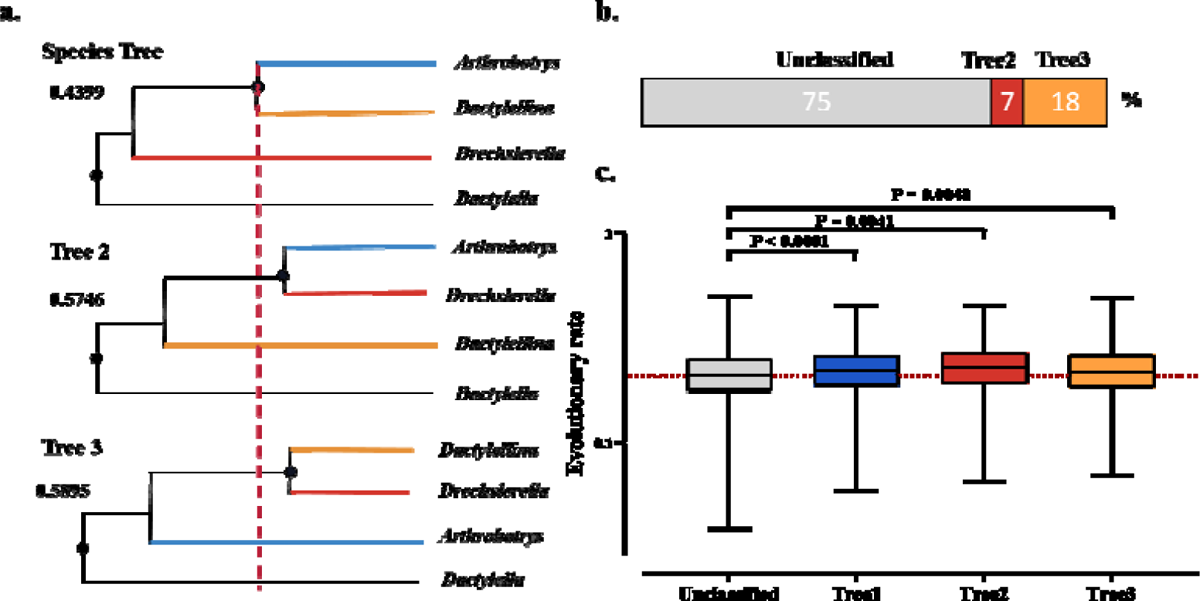
Divergence nodes and cumulative branch lengths for the three NTF genera. a. Topological structures of the three gene trees and their divergent branch lengths. b. Stacked bar chart showing the proportions of the three types of gene tree topology inconsistent with the species tree. c. Box plot of cumulative branch lengths for four types of gene trees.

The branch lengths at the divergence nodes of the gene trees likely affected by ILS were longer than those in the species tree, a significant signal supporting ILS (Song et al. 2023). We compared the divergent branch lengths between the ancestral node (node1) and the next divergence node (node2), which represents the duration of nematode-trapping device divergence in the three different types of gene trees (Figure 4a). The mean divergent branch lengths for Tree2 and Tree3 (0.5746 and 0.5895, respectively) were significantly shorter than that for the species tree (0.4399, p < 0.0001), supporting the contribution of ILS to the divergence of the three NTF lineages.

The phylogenetic conflicts in those categorized as “Unclassified” (1,036 trees) were likely caused by ILS. The MSC analysis indicated that 84.44% of the conflicts in the four lineages could not reject the hypothesis that they arose from ILS (Figure S2). ILS events involve random fixation of ancestral sequences, leading to a plethora of topologies spanning the NTF lineages. A substantial number of gene trees exhibiting inconsistency with the lineages may be due to the stochastic nature of ILS. At the same time, the lack of correspondence between these gene trees and the branches of the lineage suggest that these are more ancient ILS events, and the gene sorting may have occurred before the lineage divergence. Compared to Tree1, Tree2, and Tree3, the Unclassified type trees have significantly shorter cumulative branch lengths (Figure 4c), suggesting lower evolutionary rates (Steenwyk et al. 2021) (Supplementary Table S3).

### ILS genes under positive selection are broadly associated with growth and trap morphogenesis

Natural selection during rapid evolutionary radiation frequently leads to accelerated gene evolution and resulting phenotypic changes (Nevado et al. 2019; Hines and Rahman 2019). Positive selection among ILS genes was detected using CodeML with the site model. Sixteen single-copy orthologous genes exhibited signs of significant positive selection (Supplementary Table S2) and were enriched in functions related to the cell membrane system and cellular polarity division (Figure 5). For example, functions related to the cell membrane system include the nuclear outer membrane (GO:0005640), plasma membrane (GO:0005886), outer membrane (GO:0019867), and endoplasmic reticulum (GO:0005783). Meanwhile, functions related to cellular polarity division include the cellular bud tip (GO:0005934) and neck (GO:0005935), cellular bud (GO:0005933), and site of polarized growth (GO:0030427). Since the cell membrane system and polarity division are cellular bases for morphological innovation, the positively selected functions of these ILS genes are likely related to the morphogenesis of NTF trap structures.

**Figure 5.**
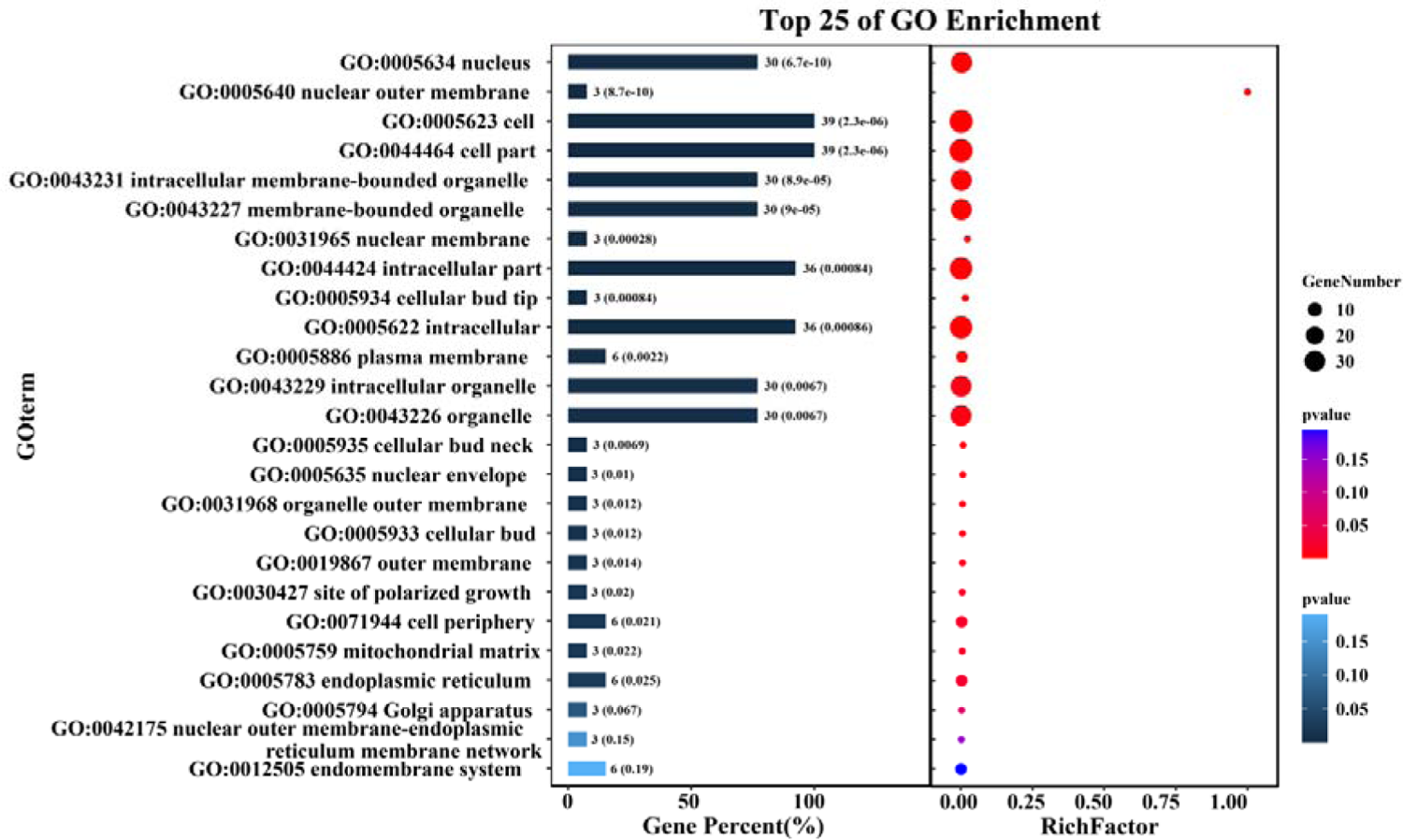
Functional enrichment analysis. Functional enrichment analysis of the ILS genes that are linked to the divergence of three NTF lineages and display signs of significant positive selection. The Gene Ontology (GO) terms enriched among those associated with the cell membrane system and polarity division are shown.

Among the conserved genes categorized as “Unclassified”, 35 gene families showed significant evidence of positive selection. The functions of these gene families are primarily enriched in processes related to the RNA polymerase, cell nucleus, and transcription (Figure S3, Supplementary Table S2). These functions are crucial for performing conserved cytological processes.

## Discussion

We investigated the evolutionary history of carnivorous NTF in Ascomycota by analyzing the genome-wide pattern of phylogenetic discordance and positive selection using the genomes of 21 species (23 strains) representing three NTF lineages. We generated the first genome-scale species tree for these NTF. While the genome-scale species tree (Figure 1) was consistent with previously published phylogenetic trees (Yang et al. 2012; Yang et al. 2007), we found extensive phylogenetic discordance across the genome and the nodes of the species tree. The ILS between lineages caused 81.3% of the phylogenetic discordance, while 18.7% were attributed to post-speciation introgression within the lineage or HGT. The reticulate phylogenetic inference indicates that introgression only led to differentiation within certain NTF genera. Although HGT events caused some conflicts between gene trees and the species tree, they are not the primary driver of the widespread phylogenetic inconsistencies. These results suggest that the PT extinction led to the rapid stochastic fixation of ancestral polymorphisms and diverged along the lineages in NTF. Subsequent positive selection accelerated the evolution of genes associated with carnivory. During this process, a small number of HGT events might have contributed to genetic polymorphism in carnivorous fungi. Moreover, gene flow between NTF lineages was restricted, with only limited introgression occurring within each lineage.

The main sources of phylogenetic discordance between gene trees and species trees are ILS, introgression, and HGT. Genome-wide signatures of ILS and introgression can be distinguished because coalescence times for regions under ILS should be older than the speciation events, whereas hybridization is the post-speciation events (Feng et al. 2022). The observation that the branch lengths from the ancestral nodes to the lineage differentiation nodes in Tree2 and Tree3 are longer than those in the species tree supports the hypothesis that ILS is the primary cause of the observed phylogenetic discordance. ILS causes ancestral genetic polymorphisms to persist during rapid speciation (Hibbins et al. 2020), and ILS events have been detected in many lineages, including marsupials (Feng et al. 2022), peat moss (Meleshko et al. 2021), butterflies (Edelman et al. 2019), and eared seals (Lopes et al. 2021). Our study indicates that the evolution of NTF represents a new case of ILS-driven evolution.

We also observed signals of introgression within *Arthrobotrys* and *Dactylellina,* with the occurrence of a reticulate phylogenetic relationship within each lineage. Consequently, the effect of introgression was more pronounced among the closely related species within the generic lineage. The role of gene introgression events in species evolution has garnered increasing attention because numerous studies have highlighted their significant effects on ecological adaptability and evolution in species such as primates, butterflies (Edelman et al. 2019), gray snub-nosed monkey (Wu et al. 2023) and foxes (L Rocha et al. 2023). Future studies should explore the effects of introgression within each NTF lineage.

Some inconsistencies between the gene trees and species tree were caused by HGT. Most HGT genes originated from bacteria, but some originated from fungi in the phylum Basidiomycota. Although HGT events may not be the main factor driving the divergence of NTF lineages, they typically introduce traits that play a crucial role in evolution (Li et al. 2022), which may also hold true for carnivorous fungi. Functional characterization of such genes should be performed to assess their significance in the evolution of NTF (Fan et al. 2024).

The most conflict-rich regions tend to be associated with the highest rates of phenotypic innovation, which have been detected in six clades of vertebrates and plants (Parins-Fukuchi et al. 2021). The most conflict-rich nodes in this study also coincide with the differentiation nodes of NTF nematode traps, which also implies that these genes undergoing ILS may be associated with morphological innovation in NTF. Here, we found that some ILS genes underwent positive selection, especially genes involved in cell membrane system have been shown to be involved in trap morphogenesis (Bai et al., 2023; Chen et al., 2022) and inflation of the constricting ring (Chen et al., 2023). The role of positive selection in the adaptive radiation of cichlids, wild tomatoes, and Jaltomata has also been demonstrated, even though there are gene tree discordances in their evolutionary process (Brawand et al. 2014; Pease et al. 2016; Wu et al. 2018). The use of gene trees for each ILS gene instead of the species tree in our positive selection analysis helped to reduce the risk of false positives. This underscores the significance of positive selection as an evolutionary driver that accelerates the adaptive radiation of carnivorous fungi.

Gene tree discordance represents another source of substitution rate variation that can lead to false inferences regarding positive selection (Mendes and Hahn 2016). Genes linked to adaptive traits might not align with the species tree, causing changes in substitution rates and potentially misleading conclusions about positive selection. Therefore, interpretation of positive selection and adaptive radiation requires caution. Our study detected positive selection in the genes associated with carnivorous traits. The use of gene trees for each ILS gene instead of the species tree in our positive selection analysis helped to reduce the risk of false positives. This underscores the significance of positive selection as an evolutionary driver that accelerates the adaptive radiation of carnivorous fungi.

Additionally, many genes that did not align with their corresponding generic clades were detected and are likely to have originated prior to the divergence of the three NTF lineages. ILS typically results in the random retention of ancestral sequences (Korstian et al. 2022; Rivas-González et al. 2023), and this stochastic process is responsible for the generation of gene trees that do not align with the clades of the lineage. The significantly shorter cumulative branch lengths observed in these gene trees (Figure 4c) suggest their ancient origin and conservation, indicating their role in conserved functions related to basic life processes rather than those associated with carnivorous lifestyle. Our findings highlight, for the first time, the importance of these genes.

## Conclusion

Through phylogenomic analyses, the evolutionary history of NTF in Ascomycota, a phylum to which most known carnivorous fungi belong, was investigated. Their evolution was facilitated by the PT extinction, which led to rapid radiation driven by ILS, coupled with positive selection of the genes associated with various carnivorous traits between generic lineages, and introgression within each lineage of two genera that form adhesive traps. These analyses advanced our understanding of the genetic mechanism underlying fungal adaptive radiation and evolution.

## Methods

### Genome mining

Genomes with published protein-coding gene predictions were obtained from the National Center for Biotechnology Information (NCBI, https://www.ncbi.nlm.nih.gov/bioproject/791178). Considering the frequent gene family expansion during fungal evolution, only single-copy genes present in all species, with cutoffs of >70% identity and >90% coverage for their cDNAs, were used in this study. In total, 2,944 gene groups were identified (Table S2, S3) using OrthoFinder v 2.5.6 (Emms and Kelly 2019). The nucleotide and protein sequences of these genes were matched. Conserved protein domains were predicted using pfam-scan (Mistry et al. 2021). Gene Ontology (GO) terms based on the functional domains were obtained using pfam2go (http://geneontology.org/external2go/pfam2go). Detailed gene functions were predicted using InterPro Scan (http://www.ebi.ac.uk/interpro/).

### Phylogenetic analyses

To minimize the impact of phylogenetic inference errors on subsequent analyses, we employed two methodologies for phylogenetic analysis, resulting in two sets of species and gene tree datasets.

The first approach involved aligning the nucleotide sequences of all single-copy orthologous genes using MAFFT v 7.520 and Gblock v 0.91b (Castresana 2000; Katoh and Standley 2013). The combined sequences were used to construct species trees using IQ-TREE v 2.2.2.7 with 1,000 replicates (Minh et al. 2020). Individual gene trees based on nucleotide sequences were constructed using IQ-TREE v 2.2.2.7 with 1,000 replicates. The species and gene trees were rooted using the corresponding sequences of *D. cylindrospora*.

The second approach involved aligning the nucleotide sequences of all single-copy orthologous genes using Clustal-Omega v 1.2.4 and ClipKIT v 2.2.2 (Sievers and Higgins 2018; Steenwyk et al. 2020b). The combined sequences were used to construct species trees using IQ-TREE v 2.2.2.6 with 1,000 replicates (Minh et al. 2020). Individual gene trees based on nucleotide sequences were constructed using IQ-TREE with 1,000 replicates. The species and gene trees were rooted using the corresponding sequences of *D. cylindrospora*.

Tree types were identified using classify_tree.py, a tool available on GitHub (https://github.com/dengweihx/classifytree). By comparing the classification results of the two datasets, only gene trees consistent across both datasets were used for further analysis.

The incongruence coefficients of gene trees at each branch node of the species tree were calculated using IQ-TREE v 2.2.2.6, and the species tree was presented using Interactive Tree Of Life (iTOL) v5 (Letunic and Bork, 2021). Densitree plots of conflicting gene tree topologies were drawn using DensiTree v 3.0.2. (https://www.cs.auckland.ac.nz/~remco/DensiTree/download.html). Pairwise Robinson-Foulds (RF) distances between gene trees were calculated using the ape package for R 4.1.3, and the RF distances were then analyzed and plotted by multidimensional scaling (MDS) (Duchene et al. 2018; R Core Team 2023). The evolutionary rate of each gene tree was calculated using PhyKIT (Steenwyk et al. 2021).

### Incomplete lineage sorting analysis

ILS signals were detected by calculating the branch lengths of the differentiated nodes of the gene trees using the Internal Branch Statistics feature of the PhyKIT toolkit (Steenwyk et al. 2021). Differences in branch length were determined using the t-test. D values were detected by z-test against whole genome backgrounds (see https://github.com/simonhmartin/tutorials/tree/master/ABBA_BABA_whole_genome for D statistics). ILS analyses based on the four-taxon branch length chi-square test were performed and plotted using the MSCquartets package R 4.1.3 (Rhodes et al. 2021). Reticulated phylogenetic inference based on the InferNetwork_MP model was performed using PhyloNet v 3.8.2 (https://phylogenomics.rice.edu/html/tutorials.html). Detection of genome-wide HGT events was performed using HGTector2 (https://github.com/qiyunlab/HGTector).

### Analysis of positive selection

Positive selection on 2,944 single-copy orthologous genes was evaluated using CodeML and PAML (Yang 2007) based on the GWideCodeML package for Python 3.10.12 (https://github.com/lauguma/gwidecodeml). The dn/ds values for each clade were calculated using the site model. To correct for errors in substitution rate estimation due to ILS, we performed branch site model calculations for the genes subjected to ILS based on their gene trees. Results from the GO term enrichment analysis were presented using the clusterProfiler package for R 4.3.2 (R Core Team 2023).

## Supporting information

Supplementary informations

## Acknowledgement

We are deeply grateful to Dr. Antonis Rokas, Department of Biological Sciences at Vanderbilt University, and Dr. Yafei Mao, Bio-X institutes, Shanghai Jiaotong University, for providing insightful advice. This work was supported by a Major International Joint Research Project grant from the Natural Scientific Foundation of China (Grant no. 32020103001) and the Startup Fund from the Nankai University to XZL. SK acknowledges support from the USDA-NIFA and Hatch Appropriation (PEN4839). JLS is a Howard Hughes Medical Institute Awardee of the Life Sciences Research Foundation.

