## Supplementary informations for "Characterization of Genome-wide Phylogenetic Conflict Uncovers Evolutionary Modes of Carnivorous Fungi"

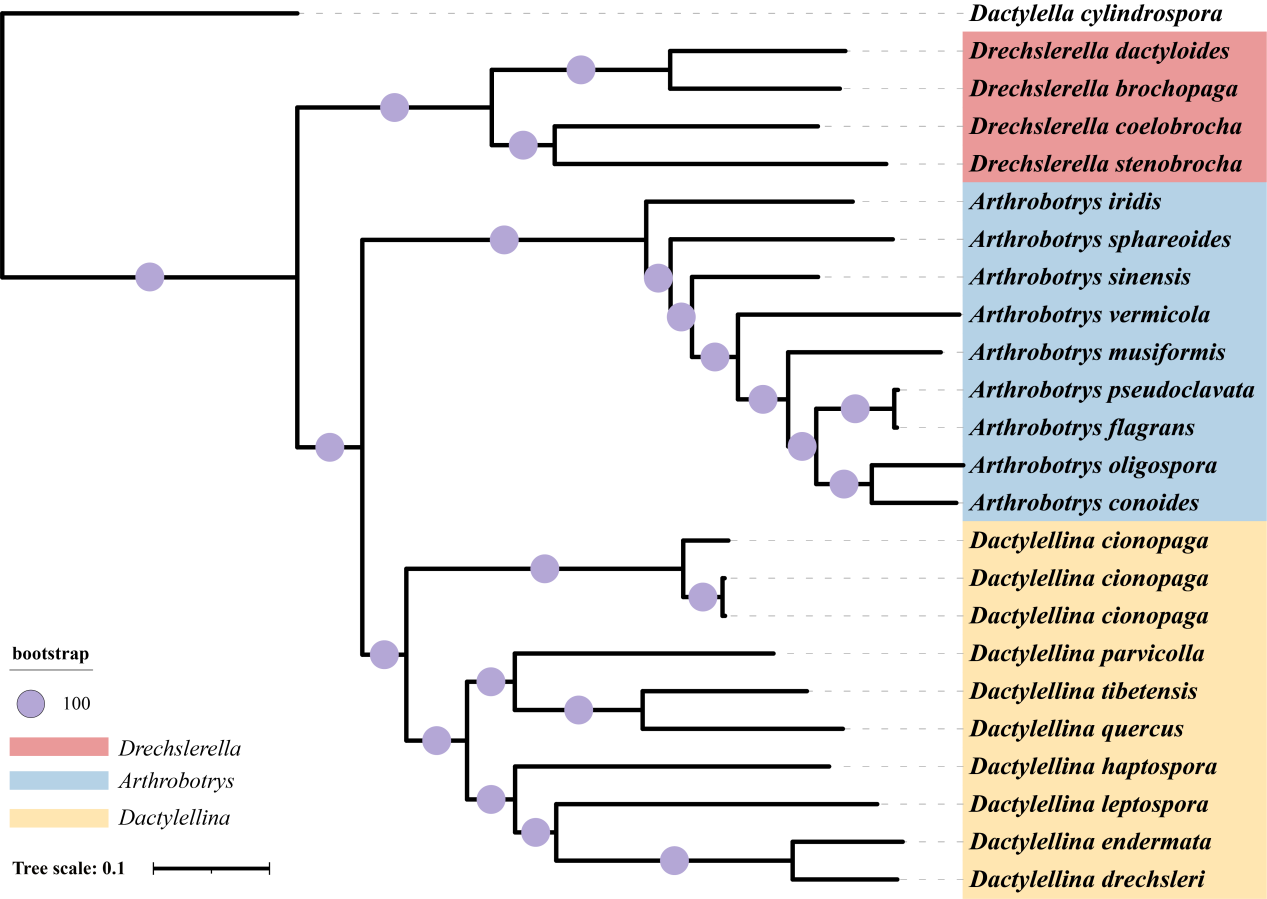

**Figure S1. Species tree2 of nematode-trapping fungi.** Their phylogenetic relationships were determined using concatenated nucleotide sequences of the single-copy orthologous genes, aligned with MAFFT, trimmed with Gblock, and a maximum likelihood tree was constructed using IQ-TREE. Bootstrap values were 100% on each node. *Dactylella cylindrospora*, a non-NTF species, was used as the outgroup.

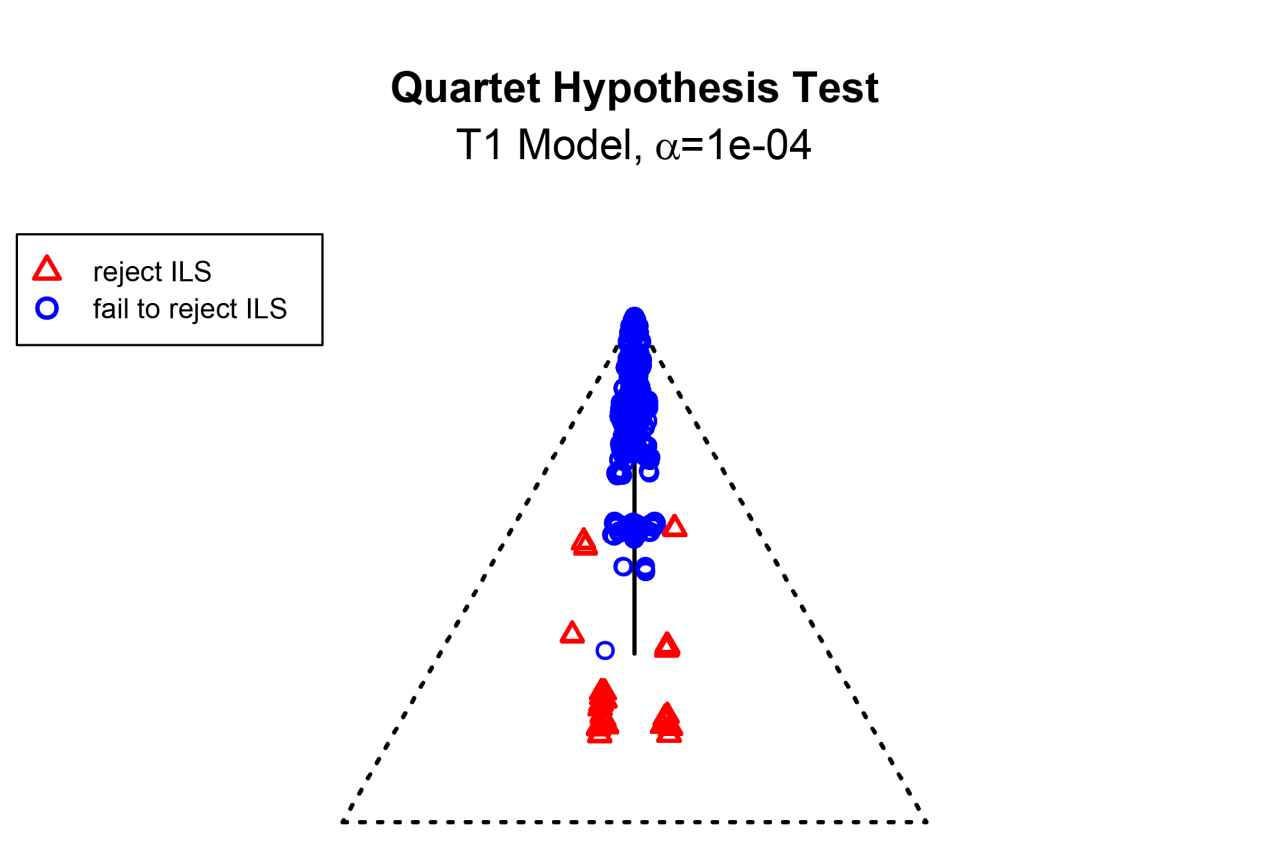
**Figure S2.** Multispecies coalescent (MSC) analysis of "Unclassified" type gene trees to evaluate the involvement of ILS. Blue circles represent four-taxa scenarios in which the topology can be explained solely by the ILS. Red triangles represent scenarios in which this hypothesis is rejected, indicating that the topology is explained by other factors. The closer the blue circles to the center of the triangle, the stronger the influence of ILS.

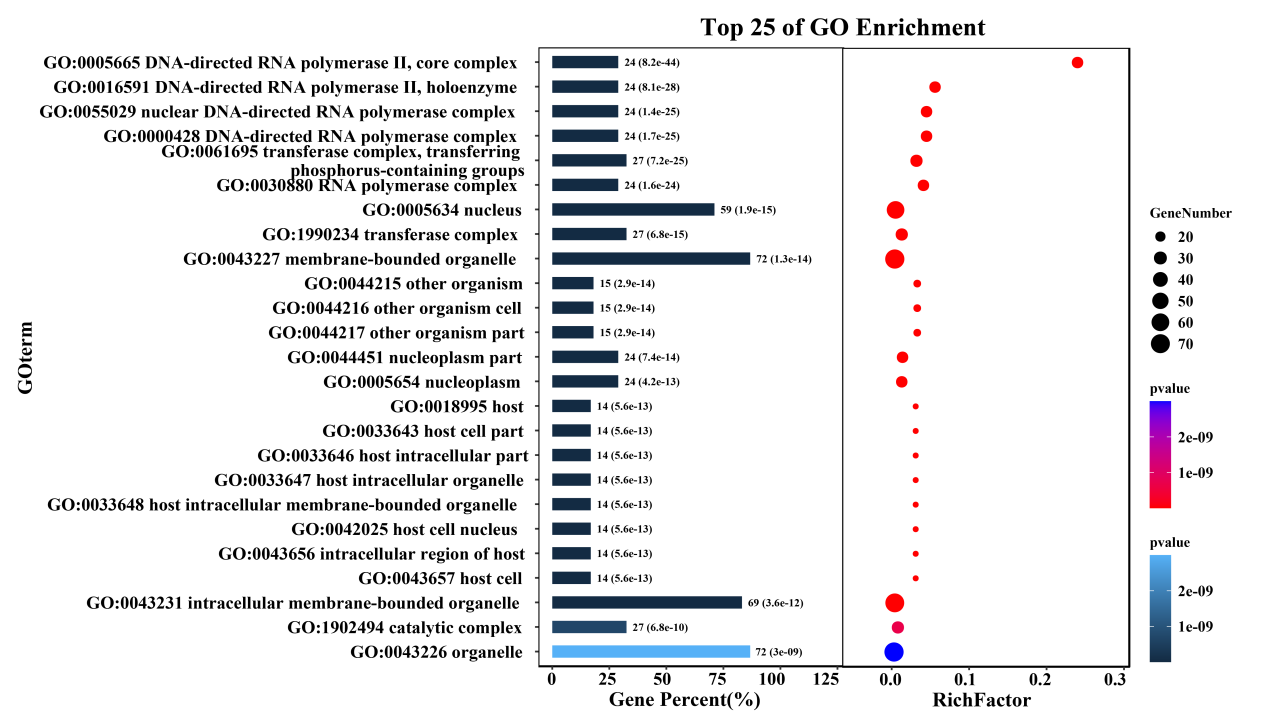

**Figure S3.** GO Functional enrichment analysis of the genes that belong to "Unclassified" Type trees.
